# Cellular Senescence is a Double-Edged Sword in Regulating Aged Immune Responses to Influenza

**DOI:** 10.1101/2023.04.10.536027

**Authors:** Blake L. Torrance, Hunter A. Panier, Andreia N. Cadar, Dominique E. Martin, Erica C. Lorenzo, Evan R. Jellison, Jenna M. Bartley, Laura Haynes

## Abstract

Clearance of senescent cells has demonstrated therapeutic potential in the context of chronic age-related diseases. Little is known, however, how clearing senescent cells affects the ability to respond to an acute infection and form quality immunological memory. We aimed to probe the effects of clearing senescent cells in aged mice on the immune response to influenza (flu) infection. We utilized a p16 trimodality reporter mouse model (p16-3MR) to allow for identification and selective deletion of p16-expressing senescent cells upon administration of ganciclovir (GCV). While p16-expressing senescent cells may exacerbate dysfunctional responses to a primary infection, our data suggest they may play a role in fostering memory cell generation. We demonstrate that although deletion of p16-expressing cells enhanced viral clearance, this also severely limited antibody production in the lungs of flu-infected aged mice. 30 days later, there were fewer flu-specific CD8 memory T cells and lower levels of flu-specific antibodies in the lungs of GCV treated mice. GCV treated mice were unable to mount an optimal memory response and demonstrated increased viral load following a heterosubtypic challenge. These results suggest that targeting senescent cells may potentiate primary responses while limiting the ability to form durable and protective immune memory with age.

## Introduction

It is well appreciated that aging results in various changes in the immune system that leave older adults at higher risk for severe infection. This is demonstrated by the fact that older adults bear the greatest burden of adverse outcomes following both COVID-19 and influenza (flu) infection (1, 2). The effects of age on the immune system are multifaceted and affect both the innate (3) and adaptive compartments (4, 5). Both CD4 and CD8 T cell responses are also diminished with age. Following flu infection, aged mice, similar to older adults, have delayed viral clearance, reduced antibody responses, and a slower resolution of inflammation following pathogen clearance (6). The mechanisms of T cell aging include both cell intrinsic and cell extrinsic factors (5). Our group and others have described that the aged microenvironment is one of the key drivers of dysfunction in CD4 and CD8 T cell responses. Adoptive transfer of aged T cells into young hosts can rescue some of the age-related changes such as reduced proliferative capacity, reduced cytotoxicity, and dysfunctional CD4 subset differentiation (7, 8). Furthermore, with age, the capacity of T cells to effectively differentiate into protective memory cells is also compromised (9). Since CD8 T cells are the key mediators of viral clearance, it is of utmost importance to understand how the aged microenvironment shapes their function, including the generation of protective T cell memory.

The aged microenvironment is characterized by a systemic increase in basal, sterile, and chronic inflammation, a state termed inflammaging (10). More recently, a role for senescent cells has emerged as a key factor in systemic increases in inflammation with age. Cellular senescence is a mostly irreversible state of cell cycle arrest that plays a key role during embryonic development and as a tumor suppression mechanism (11). Senescent cells are also required for optimal wound healing (12). After performing these physiological functions, senescent cells are typically cleared by the immune system. With age, however, these senescent cells accumulate, become resistant to apoptosis, and evade immune clearance. As these cells accumulate, they take on a heterogenous pro-inflammatory secretory program termed the senescence associated secretory phenotype (SASP) (13). SASP also has the ability to induce senescence in nearby cells highlighting their pervasive nature leading to serious detrimental health outcomes (14). Senescent cells have been linked to a wide variety of chronic diseases of aging in pre-clinical mouse models. Osteoporosis, non-alcoholic fatty liver disease, type 2 diabetes, and cardiovascular diseases have all been shown to improve following administration of senolytics, drugs that target senescent cells (15–18). However, very few studies have focused on the role of senescent cells in exacerbating age-related changes in the immune system. Work by our group demonstrated that senolytic treatment is able to improve CD4 T cell subset balance following flu infection in aged mice (19), while others have shown that senolytics improve primary responses to a mouse coronavirus (20).

The mechanistic link between senescent cells and T cell function remains unclear. Anti-viral T cell responses require tight regulation to achieve robust proliferation upon antigen encounter, engagement of strong effector phenotypes to mediate viral clearance, development of memory precursors that differentiate into protective memory cells, and regulatory signals to mediate healing and return to homeostasis following viral clearance. Senescent cells and SASP may play pleiotropic roles, restraining aspects like initial effector function while potentiating others such as memory cell precursor differentiation. We aimed to investigate this by utilizing a powerful transgenic mouse model, the p16 Trimodality Reporter (p16-3MR), which allows for identification of cells expressing p16^INK4a^ (p16), a key biomarker of senescent cells (21), as well as selective deletion of these cells to understand how cellular senescence shapes immune responses with age.

## Methods

### Mice

All experiments utilized aged (18-20mo) p16-3MR mice (12) bred and housed at UConn Health. Between 4-8 aged mice per treatment group were used. All mice underwent gross examination at the time of sacrifice and animals with obvious pathology (e.g., tumors) were excluded from the study. Original breeding pairs were generously provided by Dr. Judith Campisi. All mice were housed in a climate-controlled environment and fed standard chow and water ad libitum. All mice were cared for in accordance with the recommendations in the Guide for the Care and Use of Laboratory Animals of the National Institutes of Health. All procedures were approved by the UConn Health IACUC.

### Ganciclovir (GCV) Treatment

Mice were given 25mg/kg/day ganciclovir (GCV, Acros Organics) dissolved in PBS via intraperitoneal (i.p.) injection. Control mice were injected with equal volume PBS. Mice were treated for five consecutive days as described (12). Prior to any infection, a five day wait period was observed to ensure full clearance of GCV. Pharmacokinetic studies have demonstrated that, upon i.p. administration, systemic GCV concentrations peak 1 hour after injection and are undetectable by 2 hours (22).

### Viral Infection

Mice were placed under anesthesia with isoflurane and intranasally delivered sublethal doses of either H1N1 influenza virus A/Puerto Rico/8/34 (PR8) or H3N2 influenza virus A/HKx31 (x31). Doses for PR8 ranged from 500 EID50 in 50uL PBS for primary infection experiments or 700 EID50 in 70uL of PBS for rechallenge experiments. For x31 experiments, 3000 EID50 in 50uL PBS was used. Mice were monitored regularly to assess weight loss as an indication of infection progress. Moribund mice and mice that had lost more than 30% of their original body weight were euthanized.

### Viral Quantification

Following sacrifice, lungs were immediately flash frozen in liquid nitrogen. Lung tissue was homogenized using a handheld homogenizer (Pro Scientific) and RNA was isolated via standard trizol/chloroform (Invitrogen Life Technologies and Sigma Aldrich, respectively) extraction per the manufacturer’s protocol. cDNA was synthesized using iScript cDNA synthesis kit (Bio-Rad) using the manufacturers protocol. Viral load was determined by RT-qPCR for PR8 acid polymerase (PA) gene compared to a standard curve of known PA copy numbers as we have previously published (23). This method has been shown to directly correlate with other viral quantification methods (24). The following primer and probe were used: forward primer, 5’-CGGTCCAAATTCCTGCTGA-3’; reverse primer, 5’-CATTGGGTTCCTTCCATCCA-3’; probe, 5’-6-FAM-CCAAGTCATGAAGGAGAGGGAATACCGCT-3’ (Integrated DNA Technologies).

### Tissue Processing and Flow Cytometry

Following sacrifice, lungs were mechanically and enzymatically digested (100U/mL collagenase (Gibco)) in RPMI media containing 5% fetal bovine serum. Red blood cells were lysed using ACK lysis buffer (Gibco). Spleens were mechanically digested through 70um filters and red blood cells were lysed using ACK buffer. Lymph nodes were mechanically digested through 70um filters and did not undergo red blood cell lysis.

For flow cytometry, cells were incubated with Fc block (anti-CD16/32, ThermoFisher) followed by staining with a NP311-325 IA^b^ MHC Class II tetramer or NP366-374 H-2D^b^ MHC Class I tetramer (generated by the NIH Tetramer Core Facility). Cells were subsequently stained with surface antibodies and then either fixed using 1% paraformaldehyde or permeabilized using a FoxP3/Transcription factor fixation/permeabilization kit (ThermoFisher). Samples undergoing permeabilization were then stained with intracellular antibodies. Extended antibody information can be found in Supplemental Table 1. The PE channel was always kept clear for analysis of RFP expression by p16^+^ cells. Becton Dickinson (BD) LSR II or Bio-Rad ZE5 cytometers were used and analysis was performed using FlowJo (BD).

### Antibody Quantification

To obtain bronchoalveolar lavage (BAL), at time of sacrifice, lungs were flushed with 1mL of PBS and supernatant was collected following centrifugation to exclude cells and debris. Serum was obtained from blood collected via cardiac puncture immediately postmortem. BAL and serum were serially diluted either between 2- to 3-fold for BAL or 10-fold for serum. Diluted samples were transferred to microplates coated with either whole viral particle or flu nucleoprotein (NP). Depending on the assay, either a horseradish peroxidase conjugated to an anti-IgG or anti-IgA antibody (Southern Biotech) was used. Titer was determined at highest dilution which had a measured absorbance at 490nm over mean plus standard deviation of blanks.

### Statistics

All data are presented as mean +/- standard error of the mean (SEM). Data points outside 3 standard deviations from the mean were excluded as outliers. This resulted in the exclusion of one high outlier in the PBS group in Figure 4B. Differences between groups were determined via Student’s t-test or Mann-Whitney U-Test when data was not normally distributed as indicated by the Shapiro-Wilk test. Analyses were performed using Prism 8 software (GraphPad). p-values < 0.05 were considered significant (* p<0.05, ** p<0.01).

## Results

### Effects of Targeting p16-Expressing Senescent Cells on Primary T and B cell Responses

In this study, we utilized the p16-3MR mouse model (12) to examine the effects of p16-expressing senescent cells in both the primary and memory response to influenza. These mice express three cassettes under the control of the p16 promoter: red fluorescent protein (mRFP), luciferase, and herpesvirus thymidine kinase (Fig 1A). These cassettes allow for identification of senescent cells via flow cytometry as well as selective deletion of these cells via administration of ganciclovir (GCV), a prodrug that reacts with the herpesvirus thymidine kinase to form an apoptosis-inducing nucleoside analog causing DNA chain termination. Other groups using this model have demonstrated effective clearance of p16-expressing senescent cells upon GCV administration (25). Other studies utilizing this model have identified the presence of senescent cells in the lungs of aged p16-3MR mice (26), but this has yet to be used to study the contribution of senescent cells to diminished immune responses after flu infection in aged mice. To determine whether targeting these cells could potentiate the response to flu, we treated aged (18-20 months old) p16-3MR mice intraperitoneally with either GCV or PBS daily for five days. Following a five-day rest period after the last dose to ensure full elimination of GCV, mice were infected with a sublethal dose of H1N1 flu A/Puerto Rico/8/34 (PR8) (Fig 1B). While deletion of senescent cells did not affect weight loss following infection, it did significantly enhance efficacy of viral clearance at 12 days post infection (DPI) (Fig 1C and D, respectively). Highlighting the efficacy of this model, abrogation of p16 expression persisted up to 30 days post infection with H3N2 flu A/HKx31 (x31) (Fig 1E and F).

**Figure 1.**
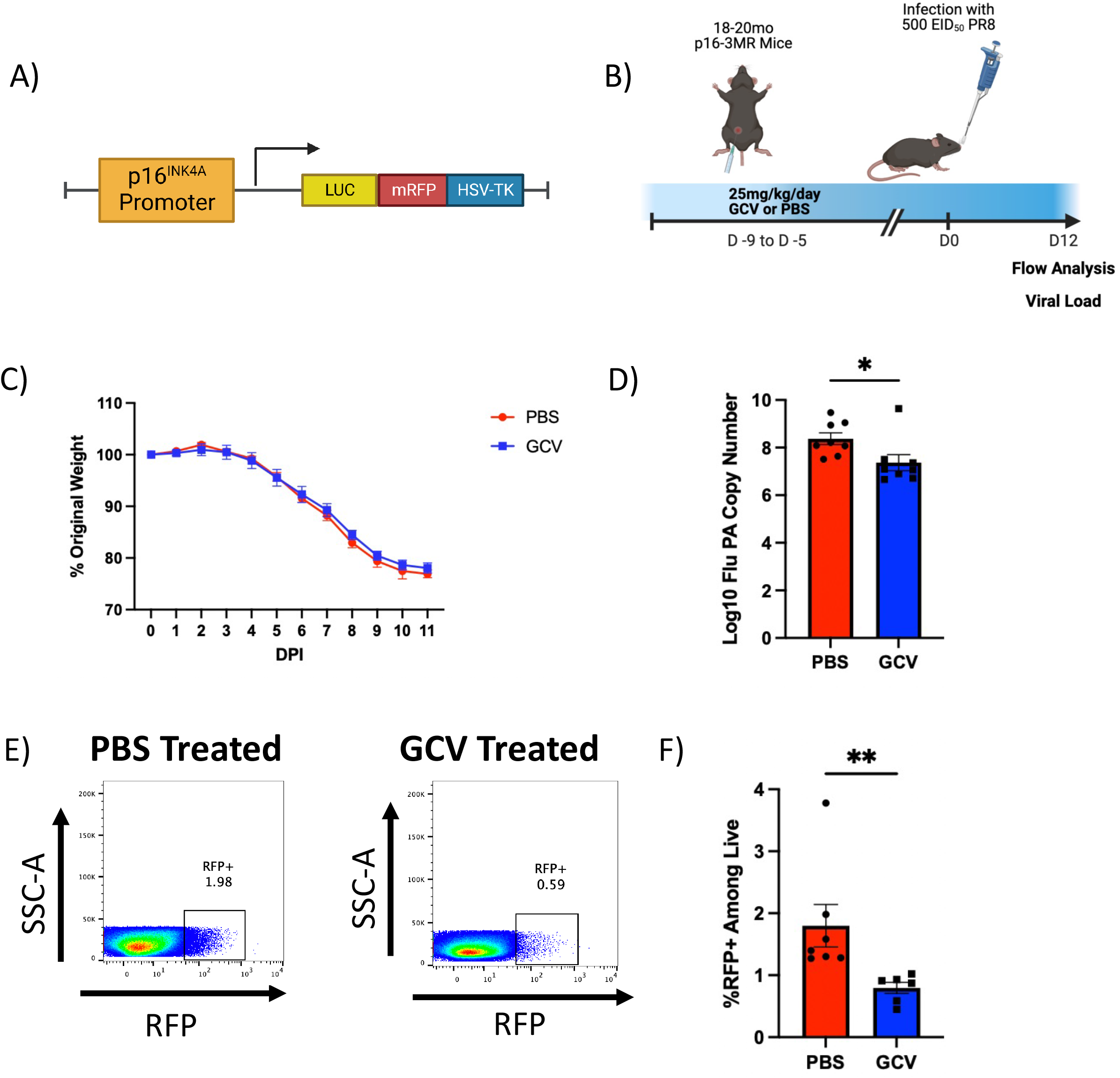
Targeting p16-Expressing Senescent Cells is Effective in Potentiating the Primary Aged Antiviral Immune Response. The p16-trimodality reporter (p16-3MR) model contains three cassettes under the control of the p16^INK4A^ promoter: one expressing luciferase, another expressing mRFP, and another expressing the herpesvirus thymidine kinase (**A**). To study the effects of p16-expressing senescent cells on the aged immune response to influenza, we treated 18-20mo p16-3MR mice with 25mg/kg/day ganciclovir (GCV) or PBS for five days. Following a 5 day rest period, we infected mice with sublethal dose of influenza (**B**). Percent of original weight lost was tracked throughout the course of infection with PR8 flu (**C**). At 12 days post infection (DPI) with PR8 flu, one cohort of mice was used to quantify viral load in the lungs via RT-qPCR (**D**). RFP expression was quantified up to 30 DPI with x31 flu to confirm efficacy of GCV treatment (**E** and **F**). Data are presented as mean +/- standard error of the mean (SEM) and each symbol represents a single animal. Mann-Whitney U-test was utilized for **D** and **F**, with a significance level of p<0.05. N=6-8 per group.

To probe the mechanism of enhanced viral clearance, we first turned to the T cell compartment. Importantly, we sought to identify the balance of short-lived effector T cells (SLECs) and memory precursor effector T cells (MPECs) via CD127 expression. CD127, or IL-7 receptor alpha, is a key marker of MPECs (27). IL-7 is an important survival signal for memory T cells, thus MPECs expressing high levels of CD127 are more likely to survive following viral clearance and become memory cells equipped for a robust secondary response upon antigen re-encounter. Lower CD127 expression is indicative of a population termed SLECs that are more effective at clearing an infection, but are more likely to die following resolution (28). While typically a minority of effector T cells, MPECs have the greatest potential to differentiate into bona fide memory cells following resolution of infection. Deletion of p16-expressing senescent cells in aged p16-3MR mice prior to flu infection resulted in fewer CD127-expressing CD8 T cells in the lungs and an overall lower mean fluorescence intensity (MFI) of CD127 expression (Fig 2A and B, Complete gating strategy can be found in Supplemental Figure 1A). We also observed that GCV treatment induced a decrease in the number and a trending decrease in the frequency of flu nucleoprotein (NP)-specific CD8 T cells infiltrating into the lung at 12 DPI (Fig 2C-E). While this may appear counterintuitive in light of the enhanced viral clearance in the GCV group, it most likely indicates that viral replication is already being well controlled and the decrease in numbers of flu-specific cells is a means to control immunopathology. Further, enhanced viral clearance results in less antigen available to continue to stimulate immune responses. Similar to the total CD8 compartment, fewer flu NP-specific CD8 T cells expressing CD127 were detected in the lungs of GCV treated mice (Fig 2F). Thus, our results show that deletion of p16-expressing senescent cells resulted in a bias toward short-lived effector CD8 T cells in the lungs and away from memory precursors, perhaps via perturbations in CD127 expression.

**Figure 2.**
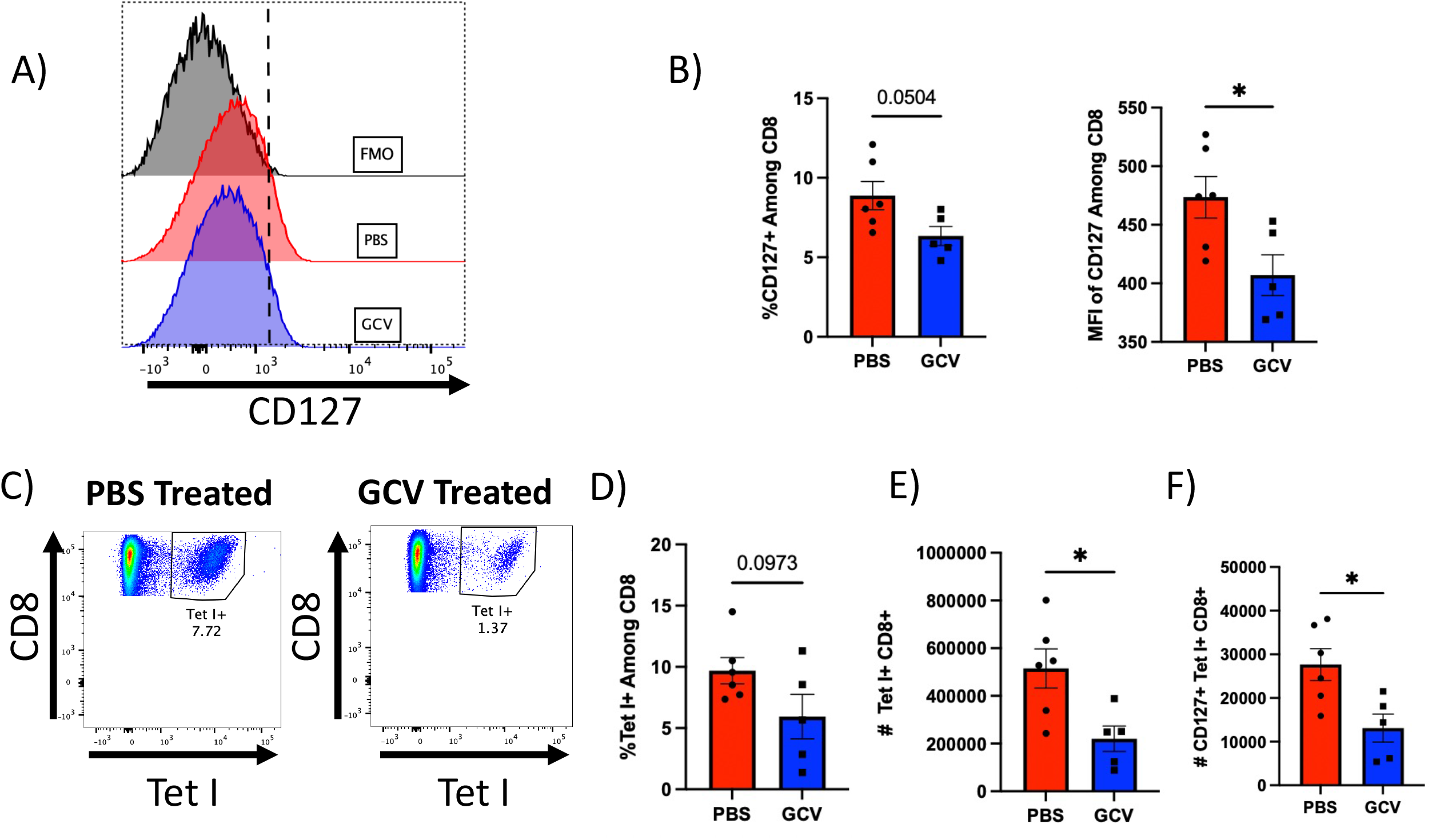
Targeting p16-Expressing Cells Decreases Memory Precursor CD8 T cells in the Lung. At 12 DPI, lung infiltrating CD8 T cells were examined for surface CD127 (IL-7Rα) expression. Both frequency of CD127+ CD8 T cells as well as mean fluorescence intensity (MFI) of CD127 among total CD8 T cells was quantified (**A** and **B**). Numbers of flu NP-specific CD8 T cells (via influenza nucleoprotein MHC class I tetramer staining) and those expressing CD127 were also quantified (**C**-**F**). All mice were infected using PR8. Data are presented as mean +/- standard error of the mean (SEM) and each symbol represents a single animal. Student’s t-tests were used for all comparisons with a significance level of p<0.05. N=5-6 per group.

Interestingly, these effects were not found among total CD4s or flu-specific CD4s (Supp. Fig 1B). In light of our previous studies examining the effects of senolytic drug treatment using a combination of dasatinib and quercetin (D+Q) on T cell differentiation following flu infection (19), we also assessed the balance of flu-specific FoxP3-expressing regulatory T cells (Tregs) and GATA3-expressing Th2 CD4 T cells and found no difference (Supp. Fig 1C). Importantly, this result with CD4 T cells highlights the differences observed when using various approaches to target senescent cells.

Examination of the B cell compartment revealed that p16 ablation did not induce any changes in class switched B cells in the lungs during flu infection (Suppl 2B). However, we found that GCV treatment induced a trending increase in the number of CD19^+^ CD138^+^ plasmablasts (Fig 3A and B). This was limited to the lungs and no similar increases in plasmablasts or CD19^-^ CD138^+^ plasma cells were observed in the spleen (Supp. Fig 2C). Despite trending increases in plasmablast numbers, virus-specific IgG production in the bronchoalveolar lavage (BAL) was found to be sharply decreased in the GCV treated groups (Fig 3C). We assayed for IgG directed towards the whole viral particle itself as well as NP, an internal flu antigen that is highly conserved across strains and can confer broad protection (29). By using the whole viral particle as a target, this approach includes quantification of a variety of neutralizing antibodies that could be directed towards any of the surface components of the particle including hemagglutinin (HA), neuraminidase (NA) and other external structures. We also assayed for IgA directed towards these targets and did not observe any differences (Fig 3C). This phenomenon was limited to local antibody production, perhaps derived from responses occurring in the bronchus associated lymphatic tissue (BALT) since no significant deficits in IgG production were found systemically (Fig 3D). Thus, deleting p16-expressing senescent cells increased the overall number of antibody-producing cells in the lung, but was associated with significant declines in the local concentration of antibody.

**Figure 3.**
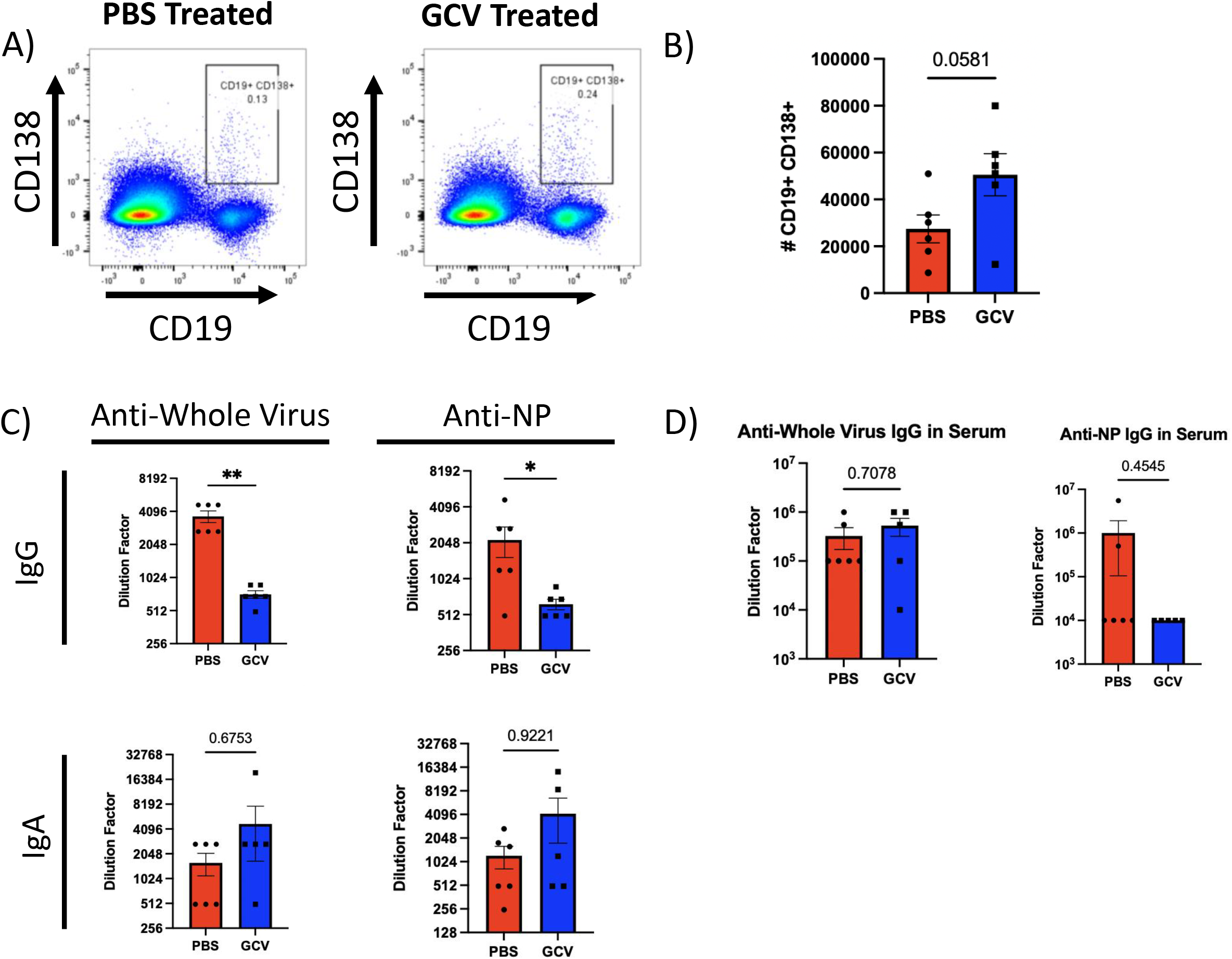
Targeting p16-Expressing Cells Alters B cell Phenotypes and Antibody Secretion in the Lung. At 14 DPI, lung infiltrating B cells were identified as plasmablasts (CD19^+^ CD138^+^) and quantified (**A** and **B**). These were initially gated on CD4 and CD8 negative cells via a dump channel (as shown in Supplemental Fig 2). Antibody titers were examined in bronchoalveolar lavage (BAL) to quantify IgA and IgG directed against either whole viral particles or flu nucleoprotein (NP) (**C**). IgG levels were quantified in the serum (**D**). All mice were infected using PR8. Data are presented as mean +/- standard error of the mean (SEM) and each symbol represents a single animal. Comparisons shown in **B** and top right panel of **C** were analyzed using Student’s t test, all other comparisons were analyzed using the Mann-Whitney U-test, all with a significance level of p<0.05. N=5-6 per group.

**Figure 4.**
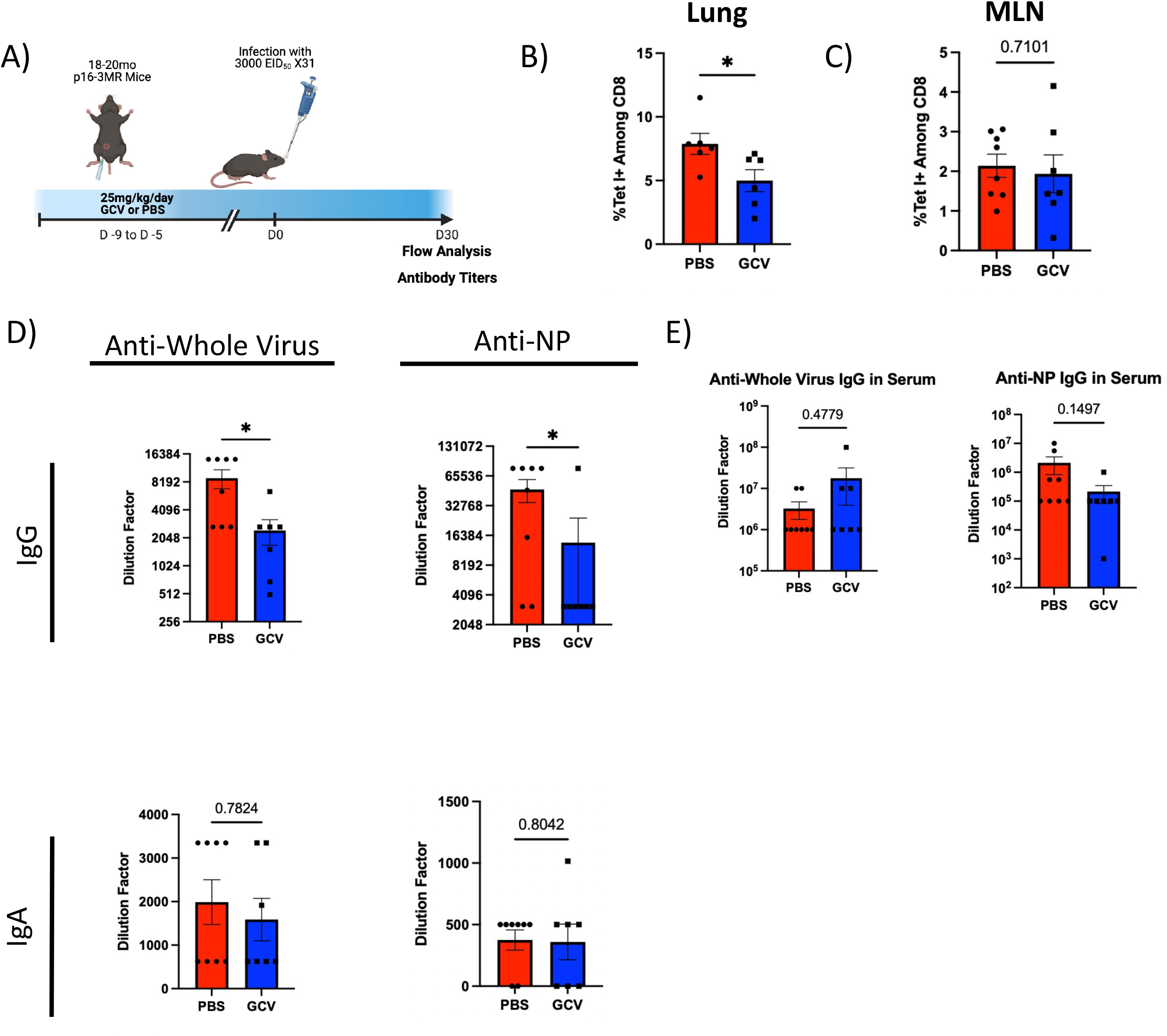
Targeting p16-Expressing Cells Limits Development of Memory CD8 T cells and Antibody Levels in the Lung. Mice were treated as in **A**. At 30 DPI, we assayed for flu NP-specific CD8 T cells remaining in the lung (**B**) and the mediastinal lymph node (MLN) (**C**). Levels of IgG and IgA directed against whole viral particles as well as flu nucleoprotein (NP) were quantified in the bronchoalveolar lavage (BAL) (**D**). Serum IgG levels against both whole viral particles and NP were also quantified in the serum (**E**). All mice were infected using x31. Data are presented as mean +/- standard error of the mean (SEM). Comparisons shown in **B** and **C** were analyzed using Student’s t-test. **D** and **E** were analyzed using the Mann-Whitney U-test, all with a significance level of p<0.05. An outlier above the mean in the PBS treated group in **B** was excluded for being more than 3 standard deviations from the mean. N=6-8 per group.

### Effects of Targeting p16-Expressing Senescent Cells on Immune Memory Formation and Function

The changes in CD127 expression that we observed during a primary infection led us to hypothesize that GCV treatment may alter memory CD8 T cell populations. To examine this, we treated aged p16-3MR mice with either GCV or PBS. Following the rest period, mice were infected with a sublethal dose of x31 as a primary infection (Fig 4A). x31 flu was chosen as the primary infection for experiments focused on memory because it is less pathogenic than PR8 and allows for later rechallenge with PR8 to test the heterologous protection conferred by memory T cells. At 30 DPI, we assessed generation of memory CD8 T cells in the lungs, since these confer the most robust protection when flu is re-encountered in the respiratory tract (30). GCV treated mice showed a marked decrease in the frequency of flu-specific memory CD8 T cells in the lungs corresponding with the observed reduction in CD127 expression (Fig 4B). No difference was observed in absolute numbers of flu-specific CD8 T cells (Supp. Fig 3A). Interestingly, no difference was observed between the two groups in the mediastinal lymph node (MLN) (Fig 4C). Specific subsets of memory cells, T effector memory (Tem, CD44^+^ CD62L^-^), T central memory (Tcm, CD44^+^ CD62L^+^), and tissue resident memory (Trm, CD103^+^ CD69^+^) were not differentially affected by GCV treatment (Supp. Fig 3B). CD4 memory T cells were also unaffected (Supp. Fig 3C). Likely, this decrease in flu-specific memory CD8 T cells is a consequence of the shift in favor of short-lived effector functions at the cost of memory precursor differentiation.

Also at 30 DPI, similar to our results at 14 DPI (Fig 3), GCV treatment resulted in sharply decreased IgG levels in the BAL against whole virus and NP (Fig 4D). Because flu NP is so highly conserved, NP-specific antibodies confer heterosubtypic cross-protection much more robustly than antibodies directed towards external viral antigens such as HA or NA, which can be highly variable across strains. As before, effects of GCV treatment on antibody concentration were limited to the site of infection and systemic IgG levels were not significantly affected (Fig 4E).

We hypothesized that while our observed deficit in memory T cell generation was moderate, the protection conferred upon pathogen reencounter would be deleteriously affected. To assess protection directly, we utilized the same GCV treatment schema as described previously (Fig 5A). In this case, at 30 DPI following x31 administration, we rechallenged with a sublethal dose of PR8 to test the protectiveness of the memory cells formed following the primary infection. Mice treated with GCV prior to the primary infection were less effective at clearing the virus compared to PBS treated controls (Fig 5B). Despite similar numbers in memory CD8 T cells in the lung (Supp. Fig 3A), the reduced protective capacity reveals a deficit in function of those memory cells formed in the absence of p16-expressing senescent cells. Therefore, our results suggest that p16-expressing senescent cells play a key role, perhaps through one or more SASP factors, in fostering the effector to memory transition in CD8 T cells.

**Figure 5.**
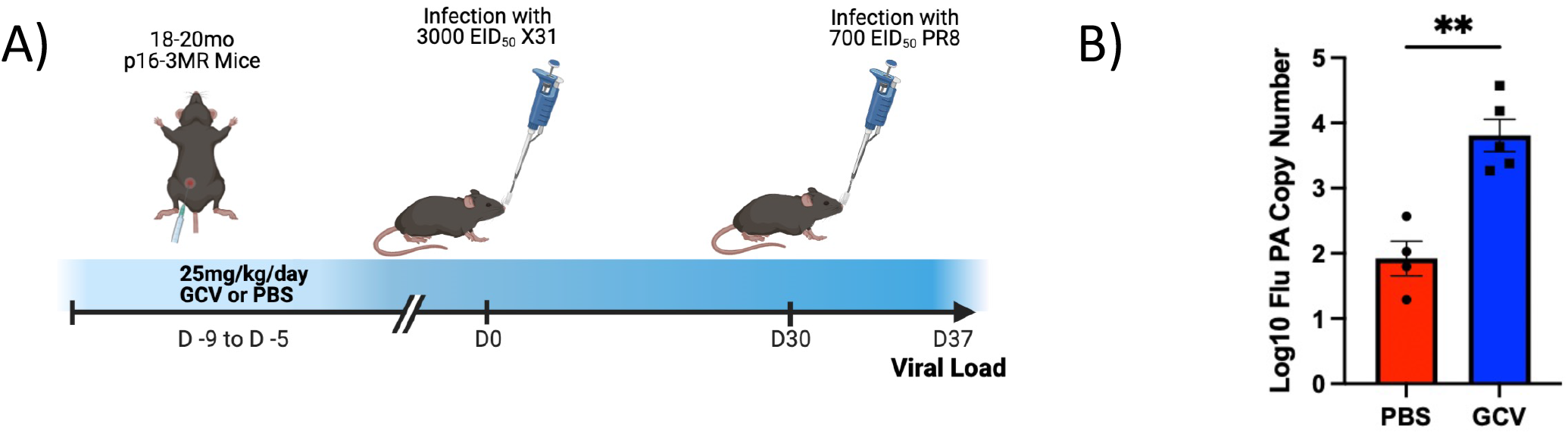
Deletion of p16-Expressing Cells Prior to Infection Impairs the Protective Function of Immunological Memory Upon Pathogen Reencounter. Mice were treated with ganciclovir or PBS and subsequently infected with x31 flu at 30 DPI, mice were rechallenged with PR8 (**A**). At 7 days following rechallenge, viral load was quantified in the lungs via RT-qPCR (**B**). Data are presented as mean +/- standard error of the mean (SEM) and each symbol represents a single animal. Comparisons were analyzed using Student’s t-test with a significance level of p<0.05. N=4-5 per group.

## Discussion

This work reveals that the role of senescent cells in aging may be more complicated than previously appreciated. Cellular senescence may be related to the altered functionality of the aged immune system, as opposed to outright dysfunction. Immune responses are complex and require a delicate balance of pro- and anti-inflammatory signals. When considering the physiological role of senescence, especially in the context of wound healing, it is possible that senescence may influence healing following an immune response. A key feature of healing is the effector to memory transition, the process by which memory precursors survive and differentiate into bona fide memory cells. Although the concept of immunological memory is foundational, the precise mechanisms by which memory cell fate is conferred are still largely unknown. Because these processes are multifaceted and involve various cell types and various soluble factors, approaches leveraging single cell-resolution are likely the best course forward. Single cell technologies have begun to unravel these processes, but much remains to be determined (31). It is unclear in our model whether the functionality of these memory cells is affected or if it is simply a quantitative effect on their development. Transcriptomic approaches may reveal important differences among these cells.

Importantly, our results here suggest that cellular senescence may be required for optimal generation of immunological memory. Work done to characterize the diversity of SLEC/MPEC CD8 T cells have indicated that early inflammatory signals can shift the subset balance between effector and memory precursors (32). Infection with different pathogens also have been shown to alter this balance via both the inflammatory environment and differences in antigen stimulation, where SLECs are driven by more persistent antigen exposure (33). It has been established that TGF-β signaling is required for optimal tissue resident memory formation and survival (34) and, importantly, TGF-β is also a common SASP factor (35). While we did not detect differences in free active TGF-β in the BAL from our experiments (data not shown), it is possible that it may not be present at time points chosen or may be at levels too low to be detected in the BAL. Interestingly, our previous work demonstrated complete ablation of TGF-β in the BAL upon D+Q treatment, albeit at an earlier time point (19). Because our results indicate deficits in both humoral and cellular memory, it is difficult to tease apart which of these are the most important in contributing to the loss of protection. Both of these mechanisms of immune memory are critically important to conferring protection and it appears that senescent cells may be involved in the development both within the site of infection.

Aside from SASP factors, it is also possible that deleting senescent cells may include structural cells such as epithelial or endothelial cells. Recently, p16-expressing fibroblasts were found to be a critical population for maintaining barrier integrity and tissue remodeling in the lungs (36). Senescent cells have also been shown to play a key role in limb regeneration in amphibian models (37). Deletion of these cells may compromise the barrier function of tissues and result in aberrant immune cell trafficking or leaking of certain cytokines or chemokines into the circulation. It is also possible that stromal cells responsible for organization of lymphoid tissues may be senescent and targeting them can exacerbate the age-related changes in organization and delineation between T and B cell zones required for optimal antibody responses. Because GCV treatment did not significantly affect numbers of antibody secreting cells while diminishing their function, perhaps senescent cells also are involved in coordinating robust antibody secretion. Further work may verify this, which would offer critical insight into the root causes of the declines in antibody production we observed in GCV treated aged p16-3MR mice.

This work stands in contrast to our own recent report utilizing senolytics prior to a primary flu infection to clear senescent cells (19). In that study, we found that D+Q administration to aged (18-20 months old) C57BL/B6 mice resulted in: (1) a significant reduction in TGF-β production in the lungs and Treg differentiation; and (2) no impact on the generation of effector CD8 T cell subsets in the lungs during the primary response. Interestingly, a study utilizing fisetin, another senolytic drug, found that targeting senescent cells improved serum antibody production (20). This highlights the heterogeneity and diversity of senescent cells and the cell types preferentially targeted by different senolytic drugs. A recent report has illustrated this by concluding that certain aging phenotypes are driven primarily by senescent cells expressing p21^Cip1^ but not p16 (38). Differences between models could help further explain discrepancies where most senolytic drugs are delivered via oral gavage and have differing pharmacokinetics than our approach delivering GCV intraperitoneally. This may result in varying degrees of senescent cell clearance and may be a confounding variable when comparing different studies.

It is also important to fully characterize both primary and secondary responses when assessing function of the adaptive immune system’s ability to combat a pathogen. The regulation of the kinetics of an immune response is dynamic and perturbations in cell populations must be probed across time, which has not yet been done in the context of targeting senescent cells to alter aged immune responses. Many outstanding questions remain in light of these results: (1) is the role of senescence in fostering the effector to memory transition relevant to a young immune system? (2) what SASP factor is responsible for altering T cell memory precursor differentiation? (3) how is the senescence environment altering the function of antibody production by B cells? In addition, our results bring to light potential concerns with the use of senolytics to potentiate immune responses in older adults. If p16-expressing senescent cells play a role in shaping the formation of durable and protective memory with age, it may be detrimental to target senescent cells with the aim to improve vaccine responses. It also is unclear how persistent the effects of clearing senescent cells are and how long it would take for senescent cells to reappear if typical aged immune responses would return as well. Clinical trials are ongoing to study effects of senolytics on COVID-19 disease severity in older adults and it will be helpful to understand how this approach may affect differentiation of memory T cells and antibody production following recovery (NCT04476953).

This, combined with further mechanistic studies in animal models, may help us to better understand cellular senescence and its effects on susceptibility to infection, generation of immunological memory, general immune function, and other processes of aging.

## Acknowledgements

The authors would like to thank Dr. Judith Campisi for the generous gift of p16-3MR breeding pairs. This study was supported by NIH grants R21AG071292 and P30AG067988 to L. Haynes. Additional support included the Diana Jacobs Kalman Scholarship for Research in the Biology of Aging awarded to B. Torrance by the American Federation for Aging Research. The authors declare no competing financial interests.

---

**Supplemental Figure 1.**
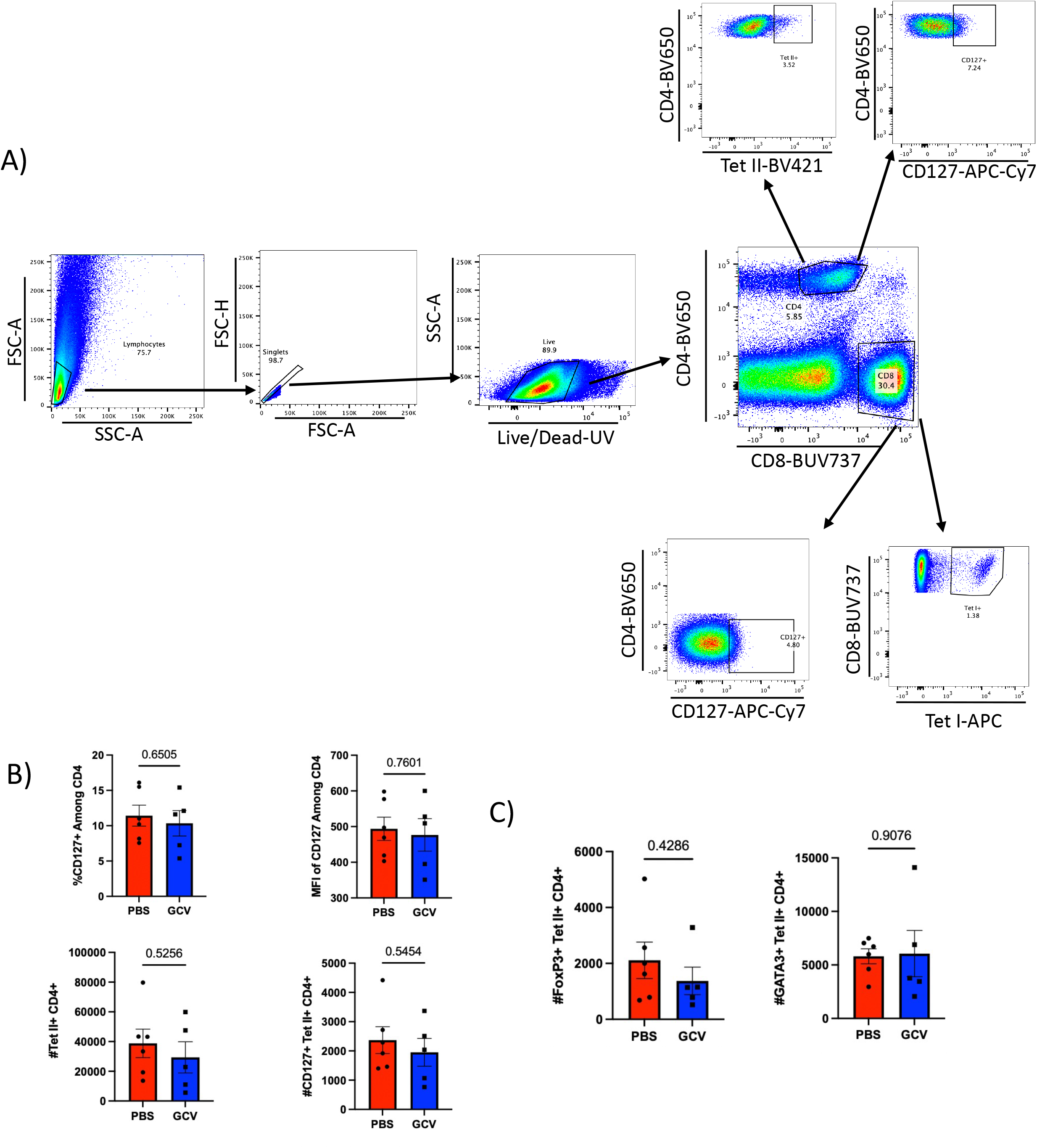
Alterations in Primary T cell Responses in the Absence of p16-Expressing Cells are CD8 Specific. All T cell phenotyping utilized the gating strategy shown in **A**. At 12 DPI, CD4 T cells were identified as memory precursors in the general CD4 and Flu NP-specific compartments (**B**). FoxP3 and GATA3 expression were quantified among Flu NP-Specific CD4s (**C**). Data are presented as mean +/- standard error of the mean (SEM) and each symbol represents a single animal. Left panel of **C** was analyzed using Mann-Whitney U-test. All other comparisons were analyzed using Student’s t-test with a significance level of p<0.05. N=5-6 per group.

**Supplemental Figure 2.**
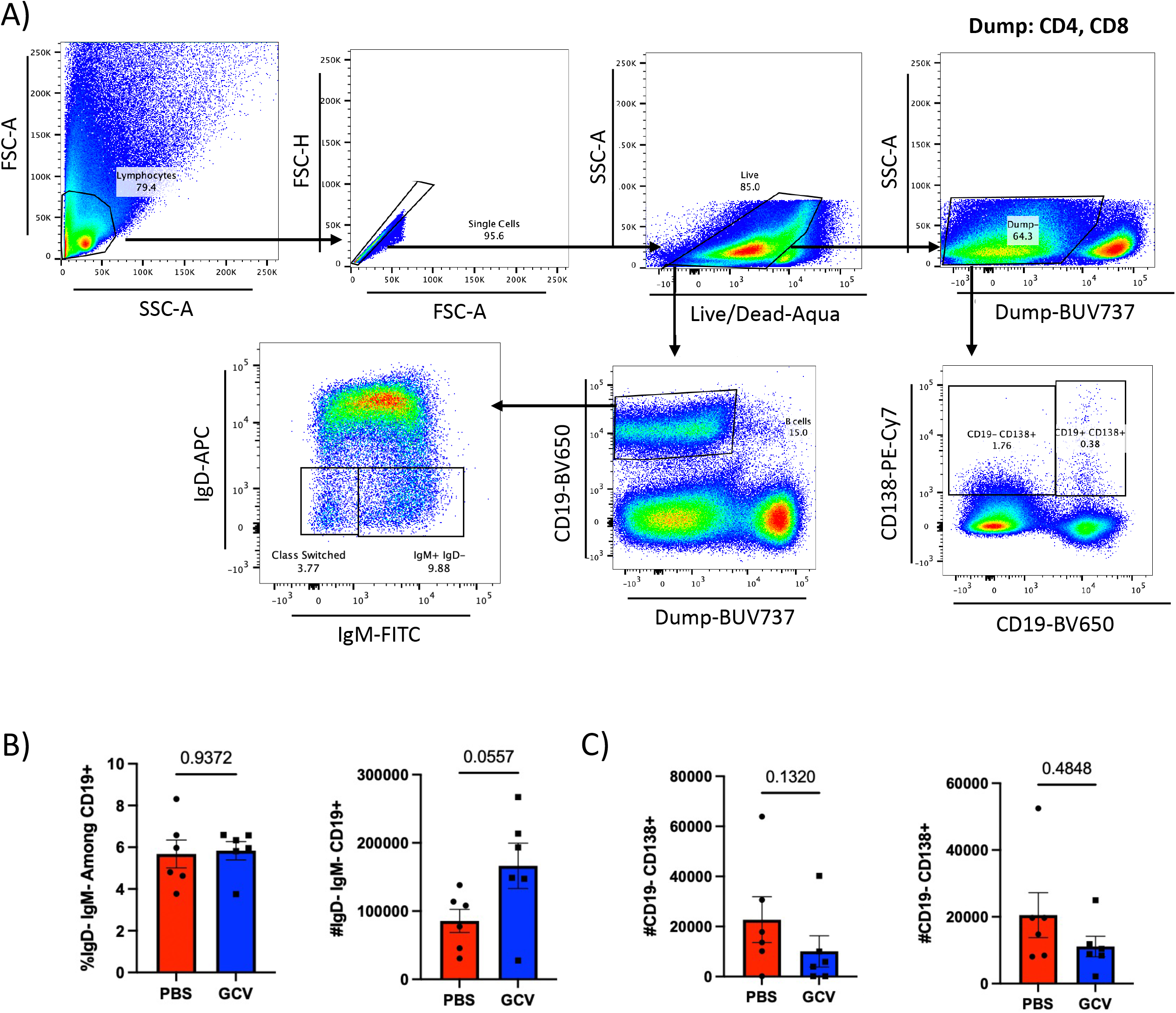
Flow Cytometric Analysis of B cell Phenotypes Following Flu Infection. All B cell phenotyping utilized the gating strategy shown in **A**. A dump channel containing anti-CD4 and CD8 was used. At 14 DPI, B cells in the lungs were assayed for expression of IgD and IgM to determine class switched status (**B**). At 14 dpi, spleen plasmablasts (CD19^+^ CD138^+^) and plasma cells (CD19^-^ CD138^+^) were quantified (**C**). Data are presented as mean +/- standard error of the mean (SEM) and each symbol represents a single animal. Comparisons in the left panel of **B** and all of **C** were analyzed using Mann-Whitney U-test. All other comparisons were analyzed using Student’s t-test with a significance level of p<0.05. N=6 per group.

**Supplemental Figure 3.**
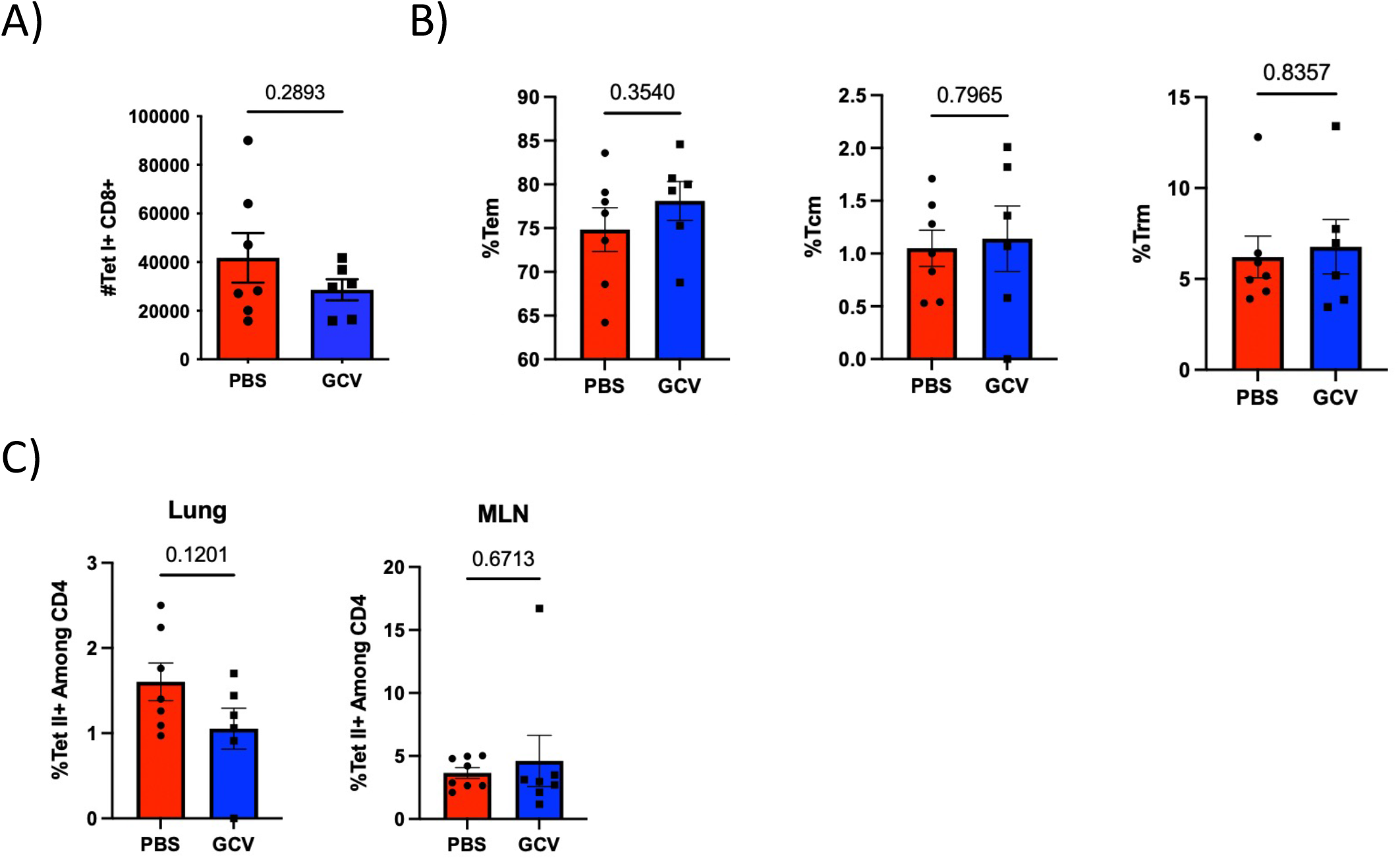
Alterations in Generation of T cell Memory in the Absence of p16-Expressing Cells are CD8 Specific and Do Not Affect Memory Subsets. Mice were infected with sublethal dose of X31 H3N2 flu. At 30 DPI, Flu NP-specific CD8 T cells in the lung were quantified (**A**) and classified into memory subsets (**B**). Frequency of Flu NP-specific CD4 T cells was also assessed in the lung and mediastinal lymph node (MLN) (**C**). Data are presented as mean +/- standard error of the mean (SEM) and each symbol represents a single animal. Comparison in the middle panel of **B** and right panel of **C** were analyzed using Mann-Whitney U test. All other comparisons were analyzed using Student’s t-test with a significance level of p<0.05. N=6-7 per group.

**Supplemental Table 1.** Extended Antibody Information for Flow Cytometry Experiments.

| **Specificity** | **Fluorochrome** | **Clone** | **Dilution** | **Vendor/Supplier** |
| --- | --- | --- | --- | --- |
| CD4 | BV650 | RM4-5 | 1:100 | BD Biosciences |
| CD4 | BUV737 | RM4-5 | 1:300 | BD Biosciences |
| CD8a | BUV737 | 53-6.7 | 1:100 or 1:300 | BD Biosciences |
| NP<sub>311-325</sub> IA<sup>b</sup> MHC Class II tetramer | BV421 | N/A | 1:50 | NIH Tetramer Core |
| NP<sub>366-374</sub> H-2D<sup>b</sup> MHC Class I tetramer | APC | N/A | 1:50 | NIH Tetramer Core |
| CD127 | APC-Cy7 | AZR34 | 1:100 | Biolegend |
| Tbet | BV711 | 4B10 | 1:100 | Biolegend |
| FoxP3 | AF700 | FJK-16s | 1:100 | ThermoFisher |
| GATA3 | FITC | 16E10A23 | 1:200 | Biolegend |
| Bcl6 | PerCPef710 | BCL-DWN | 1:1000 | ThermoFisher |
| CD44 | FITC | IM7 | 1:100 | BD Biosciences |
| CD62L | APC-Cy7 | MEL14 | 1:100 | BD Biosciences |
| CD127 | BV711 | AZR34 | 1:100 | Biolegend |
| CD69 | BV605 | H1.2F3 | 1:100 | BD Biosciences |
| CD103 | PerCP-Cy5.5 | 2E7 | 1:100 | Biolegend |
| CD19 | BV650 | 6D5 | 1:200 | Biolegend |
| CD138 | PE-Cy7 | 281-2 | 1:100 | Biolegend |
| IgD | APC | 11-26c | 1:300 | ThermoFisher |
| IgM | FITC | 11/41 | 1:300 | BD Biosciences |

